# Template switching in DNA replication can create and maintain RNA hairpins

**DOI:** 10.1101/2021.04.15.439925

**Authors:** Heli Mönttinen, Ari Löytynoja

**Affiliations:** Institute of Biotechnology, HiLIFE, P.O.Box 56, 00014 University of Helsinki, Finland

## Abstract

The evolutionary origin of ribonucleic acid (RNA) stem structures and the preservation of their base-pairing under a spontaneous and random mutation process have puzzled theoretical evolutionary biologists. DNA replication-related template switching is a mutation mechanism that creates reverse-complement copies of sequence regions within a genome by replicating briefly either along the complementary or nascent DNA strand. Depending on the relative positions and context of the four switch points, this process may produce a reverse-complement repeat capable of forming the stem of a perfect DNA hairpin, or fix the base-pairing of an existing stem. Template switching is typically thought to trigger large structural changes and its possible role in the origin and evolution of RNA genes has not been studied. Here we show that the reconstructed ancestral histories of RNA genes contain mutation patterns consistent with the DNA replication-related template switching. In addition to multi-base compensatory mutations, the mechanism can explain complex sequence changes, though mutations breaking the structure rarely get fixed in evolution. Our results suggest a solution for the longstanding dilemma of RNA gene evolution and demonstrate how template switching can both create perfect stems with a single mutation event and help maintaining the stem structure over time. Interestingly, template switching also provides an elegant explanation for the asymmetric base-pair frequencies within RNA stems.

---

The characteristic feature of RNAs are the helical structures formed by intra-molecular pairings of two to ten consecutive complementary bases (1–3). The basis of helices are the hydrogen bonds between classical Watson-Crick pairs and less stable G-U pairs, and their structures are further stabilized by the stacking interactions between successive base pairs. While the underlying sequences may evolve, the locations of RNA helices are highly conserved among related sequences. Computational methods for inference and validation of RNA secondary structures exploit this (4; 5), and correlated base changes turning one Watson-Crick pair to another, e.g., A-U *→* G-C, are considered the ultimate evidence of a functionally important stem. Such compensatory mutations (CMs) are commonly believed to happen via an intermediate state involving G-U base pairing (5). However, the rate of simultaneous substitution of both members of a base pair has been shown to be greater than zero (3; 6), suggesting that either the intermediate state with non-Watson-Crick pairing is very short or RNA stem sequences can also evolve through double substitutions. Apparent simultaneous double substitutions have been explained with population models where the initial mutations are kept at low frequency by selection and only increase in frequency after another change has restored the base pairing (3). This still requires a very high rate of matching mutations within the population.

We showed earlier that DNA replication-related template switch mutations (TSMs) (7; 8) can produce reverse-complement repeats needed for perfect DNA hairpins and fix the base-pairing of existing stems (9). The mechanism has been thoroughly studied in microbes (7; 8; 10; 11) and been aware of in eukaryotic research (12–14), but few studies have looked at the role of the resulting mutations in genes (though see (15)). Here we set out to investigate the role of TSMs in the evolution of functional genes, especially that of RNA genes dependent on stem structures formed by reverse-complement sequences. We used inferred ancestral sequences to reconstruct the mutation history for sets of closely related RNA sequences and located individual mutations into specific tree branches (Fig. 1a). Using the sequence histories, we identified *de novo* hairpins and analyzed CMs consistent with the “two-step process” (3). In the latter, the intermediate state is observed such that the initial mutation breaking the base pairing is placed in one tree branch and the compensatory change in another branch (Fig. 1a). Template switching could then either trigger a matching change in the unmutated stem or restore the original base pairing by copying the unmutated sequence. We hypothesized that if TSMs are involved in the CM process, they should occasionally leave patterns where the CM is associated with parallel changes (i.e. mutations that according to the phylogeny appeared concurrently) in its immediate proximity (Fig. 1b-f). Within RNA sequences, such associated changes would appear as multiple compensatory changes in the stem, extension of the stem region, matching mutations within the loop, or as fully inverted loops (Fig. 1f).

**Fig. 1.**
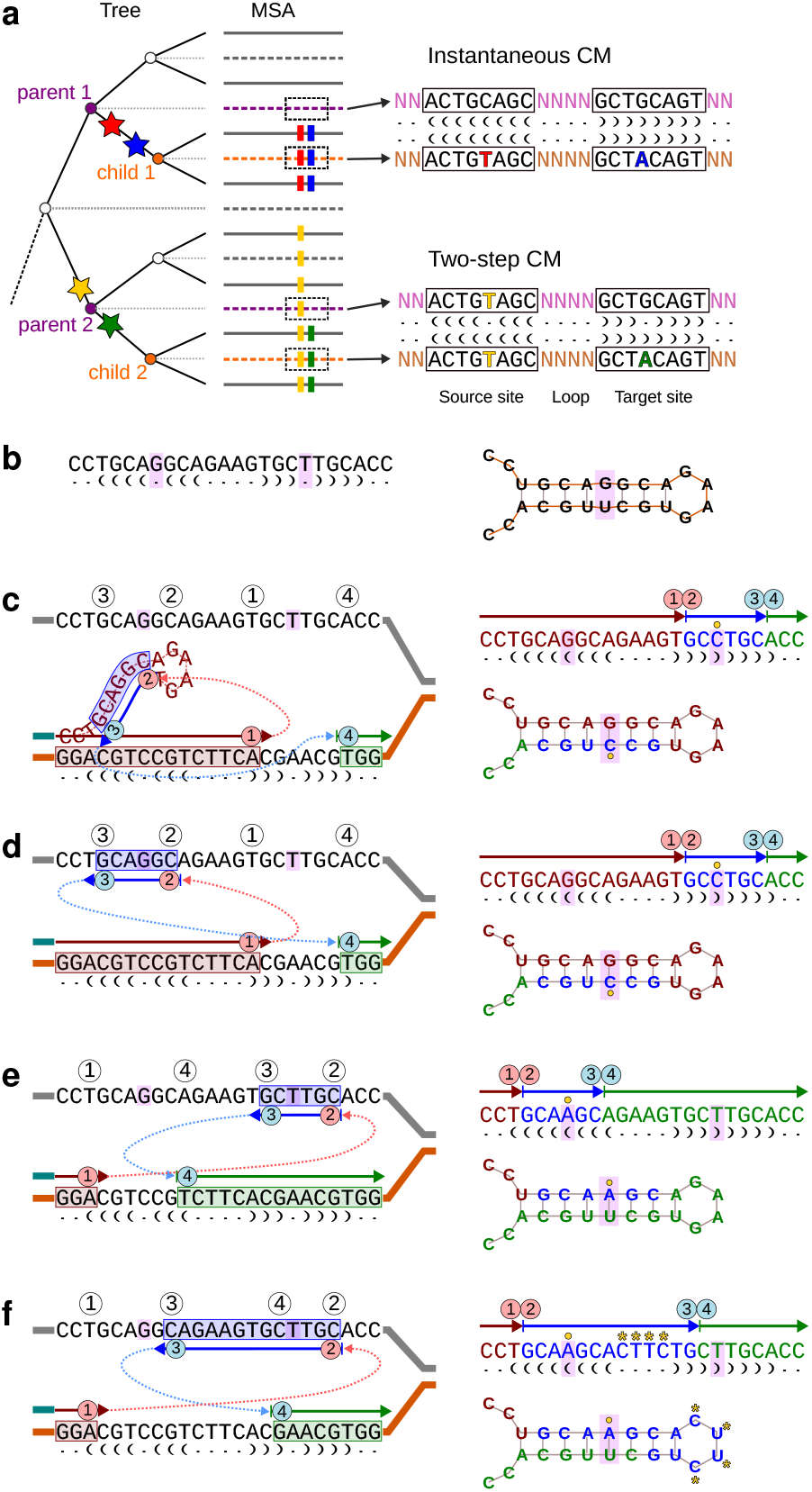
Compensatory mutations and TSM process. **a** Inference of ancestral sequences (dashed lines) for the internal nodes of a phylogenetic tree allows locating mutations (stars) to specific tree branches. Initial and restoring mutations happening in the same branch (red and blue) appear as instantaneous CMs and the perfect stem base pairing is retained. If the initial and restoring mutation appear in different branches (yellow and green), the mechanism triggering the CM can be studied. **b** A hypothetical sequence forming an RNA stem structure with a noncanonical G-U pair. **c-f** The Watson-Crick pairing can be corrected by DNA replication (solid arrows; first red) briefly switching to another template (blue arrow) and then returning to the original strand (green arrow). The outcome of the process (right) depends on the relative positions of the four switch points, ➀ - ➃. **c** In intra-strand switch, the newly synthesized sequence (in red) forms a hairpin and templates the replication. **d** In inter-strand switch, the complementary strand templates the replication. A backwards jump (➁ ¡ ➀) corrects the mismatch with a G-C pair (yellow circle) and cannot cause associated mutations within the loop. **e** Inter-strand switch forwards (➀ ¡ ➁) corrects the mismatch with an A-U pair and, depending on positions of the switch points, may cause associated mutations. When the source sequence (in blue) is within the stem region, only the mismatch is fixed. **f** Source sequence covering the loop region inverts the loop sequence, causing multiple parallel changes (yellow asterisks). Dot-bracket notations show predicted secondary structures. Opening and closing brackets indicate pairing bases in the stem, unpaired bases are marked with dots.

We tested the CM hypothesis with ribosomal RNA (rRNA) sequences that evolve at a high rate (3) and for which large quantities of data are available due to their use as phylogenetic markers (16). Ribosomal DNA (rDNA) genes usually appear as multiple copies in the genome and evolve in a concerted fashion (17). While non-homologous recombination among the copies complicates the analyses by producing conflicting phylogenetic signal, the high copy number also elevates the overall mutation rate and variants temporarily segregating within gene clusters may be detectable with DNA sequencing. In line with our hypothesis, we identified mutation patterns consistent with the TSM mechanism both among historical changes separating established evolutionary lineages and among recent mutations, likely destined for removal by drift and selection. Unexpectedly, our analyses of the stem loop sequences suggested a solution for another dilemma in the RNA evolution, the asymmetry of the base-pair substitution process in the RNA stems (3).

## Results

### Reverse-complement repeats at novel RNA stems

We analyzed mammalian genomic sequences around annotated human microRNAs and found mutation patterns consistent with the TSM mechanism (Fig. 2a-c). The phylogenetic analysis supported the creation of microRNA loci as reverse-complement repeats and occurrence of the potentially functional bulge-causing mismatches after a phase of perfect stem pairing. In the case of MIR633, the role of the TSM mechanism was strengthened by a subsequent inversion of the loop sequence in the evolutionary lineage leading to baboons (Fig. 2a,d). We found similar TSM-like patterns creating novel stems within rRNA genes (Fig. 2e-f). A notable difference to single copy microRNA genes was the phylogenetic inconsistency of the mutation patterns in the multi-copy rRNA genes, probably reflecting recombination among the non-identical gene copies. Nevertheless, the results suggested that TSM-like patterns are found in functional RNA genes.

**Fig. 2.**
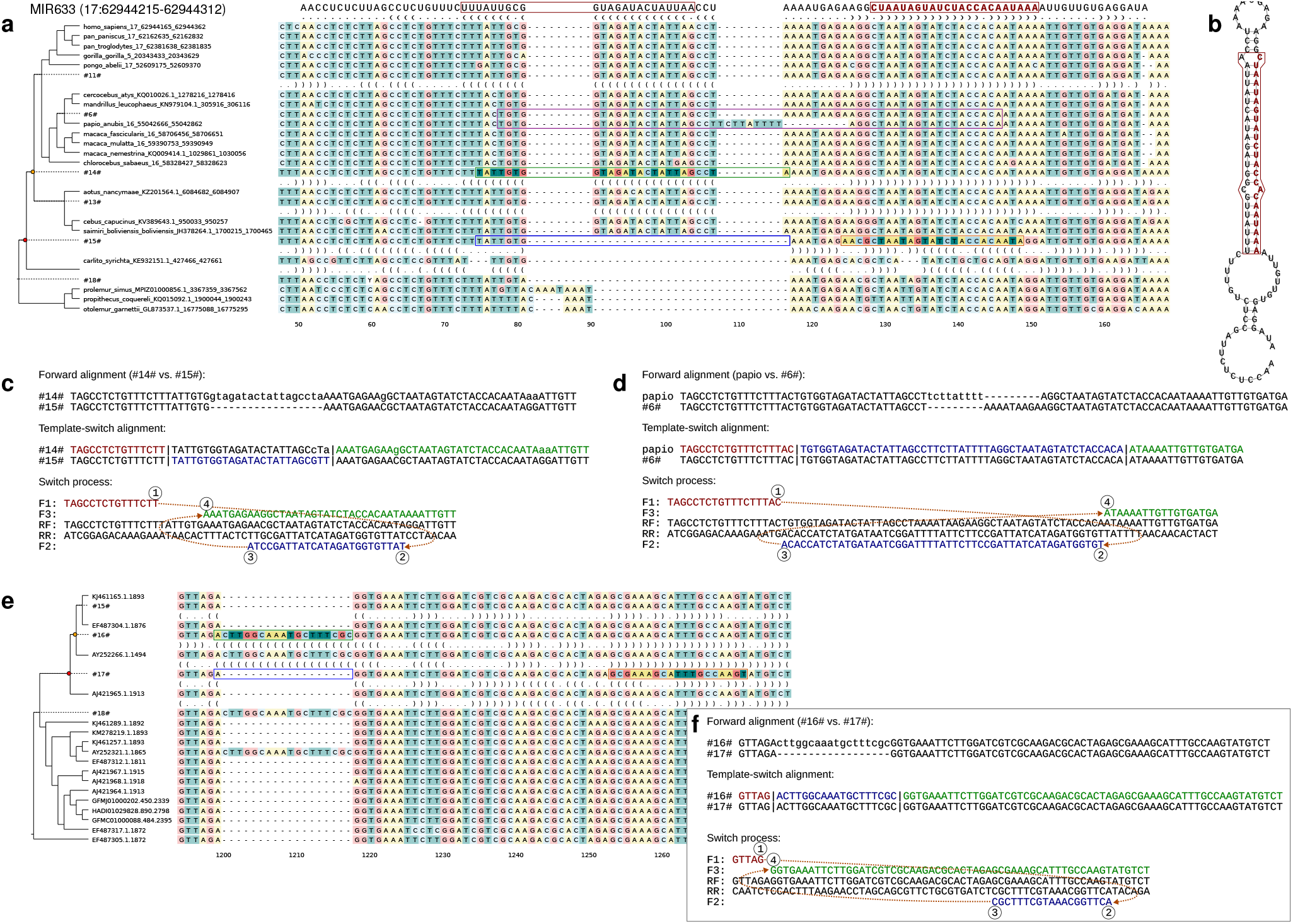
Novel stem structures consistent with template switching. **a** The MIR633 sequence is found in monkeys and apes and appears to have been created by a TSM event after the split of Tarsiiformes. The microRNA sequence is shown on top with the functional region highlighted in red. The inferred TSM has occurred between the nodes #15# and #14#: the inferred source and target loci are framed with orange and green boxes while the homologous ancestral sequence is marked with blue. **b** The inferred stem structure for MIR633. **c** The TSM process explaining the creation of the stem structure required for MIR633. **d** The differences in the olive baboon loop sequence can be explained by a subsequent TSM event. **e** An insertion changing the inferred structure of insect rRNA molecules can be explained by a TSM event between the nodes #17# and #16#. Inconsistency of the insertion pattern and the overall sequence phylogeny indicates recombination among the rRNA copies. **f** The TSM process explaining the insertion.

### Reconstruction of ancestral rRNA gene histories

To study the role of the mechanism in the CM process, we downloaded 614,502 large and small subunit rRNA sequences from Rfam database (v. 14.1) (18) and Silva rRNA database (v. 138) (16), and clustered them by minimum sequence identity of 90% and 75% pairwise coverage (Fig. S1; see Methods). To reduce the sequence numbers while retaining local dissimilarities, we used a sliding window approach and kept sequences showing at least two differences within a 20-bp window. This resulted in 5,525 clusters with at least 10 sequences, containing in total 289,024 sequences. After aligning the sequences, we inferred a maximum likelihood tree for each cluster and reconstructed ancestral sequences for the internal nodes (Fig. 1a).

Using the reconstructed sequence history, we analyzed each *parent node* - *child node* pair in the tree and identified mutation clusters between the corresponding sequences that were explainable by a TSM. We considered secondary structures for the sequences involved as well as for their immediate parent and descendant sequences (see Methods), and extracted the cases in which a CM restores the mismatch in the parent sequence to perfect base-pairing in the child sequence (Fig. 1a, “Two-step CM”). Due to the short length of rRNA stems, a statistic proposed for genomic TSM events (19) could not be applied for the evaluation of inferred events. Instead, we required a minimum length and number of sequence changes explainable by the TSM (Table 1) and applied strict criteria on sequence and structure conservation (see Methods). Given the regions passing the original quality control, we studied the statistical significance of the detected TSM-like patterns using simulations. For each cluster, we sampled one leaf sequence and simulated sequences according to the maximum likelihood tree. Using the same quality criteria as for empirical data, we computed the number of TSM-like patterns among five replicates of simulated data. Finally, we confirmed the quality of structure predictions by comparing them against a set of solved structures in Protein Data Bank. We found the locations of hairpins to be correctly inferred while the loop boundaries and the locations of bulges and internal loops contained minor error (Fig. S2), largely explained by the structure prediction method only considering the noncanonical pair G-U.

**Table 1.**
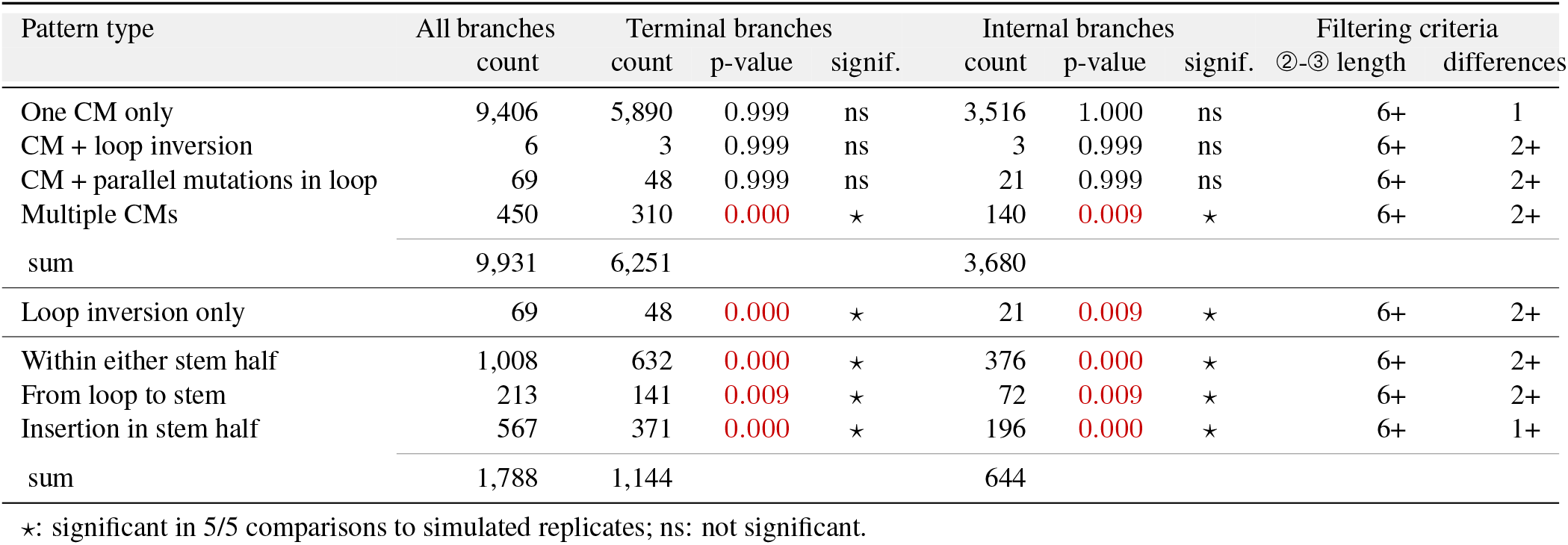
Different mutation patterns explainable by a single TSM. Types indicated by asterisks are statistically enriched in empirical data.

### Footprints of compensatory mutations

The underlying assumption of our analyses is that a great majority of the RNA stems are optimized with perfect Watson-Crick base pairing and evolution selects for congruent CMs. We found in total 9,931 cases of two-step CM to pass our quality criteria requiring ➁-➂ length of 6 bp with a maximum distance of 8 bp between the TSM source and target regions (6,251 at terminal and 3,680 at internal branches, see Table 1). Of these, 94.7% were isolated base changes and uninformative about the underlying mutation process, while the remaining 525 cases were consistent with the TSM mechanism causing a CM and a parallel change (Table 1).

We hypothesised that a CM associated with an inversion of the loop sequence (cf. Fig. 1f) would provide the strongest possible support for the model. Such cases were extremely rare (Fig. 3a) and even the CMs associated with changes in the stem loop sequence were depleted in empirical data in comparison to simulated sequences (Table 1, Table S1). The loop sequence is known to be important for the bending of the RNA strand, however, and the sequence patterns are highly conserved (20; 21). Outside the loop sequence parallel changes were more common and the cases of TSM-like patterns simultaneously converting multiple mismatches to perfect basepairing were statistically enriched in empirical data (Table 1). The enrichment was especially strong in terminal nodes which may reflect reduced accuracy of ancestral state reconstruction or the impact of negative selection in the deeper internal nodes.

**Fig. 3.**
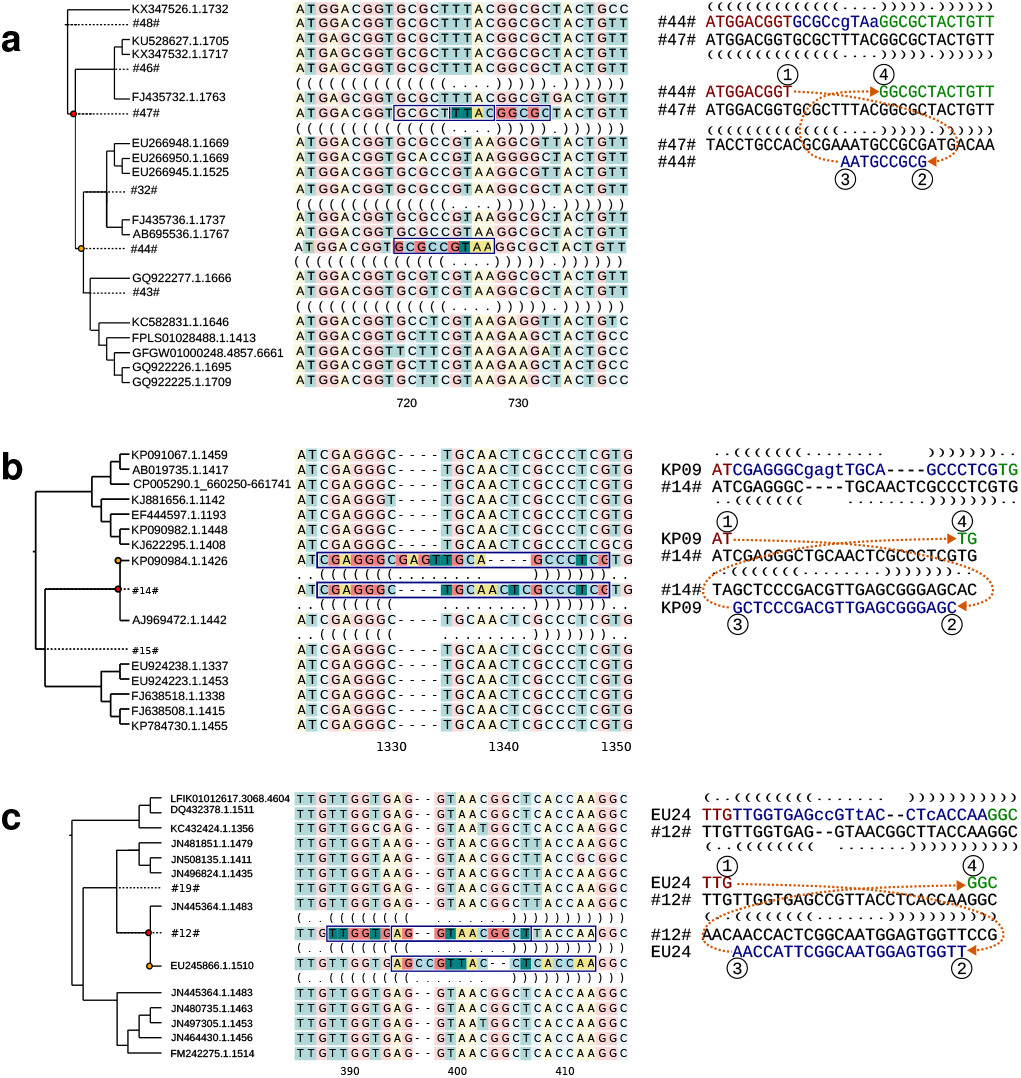
TSM-like patterns with different effects on sequence and secondary structure. Alignment shows inferred sequences for selected ancestral nodes and predicted secondary structures for the affected nodes and their immediate child nodes. The source and target regions for the inferred TSM event are highlighted and the homologous region in the parent is framed. Sequences on the right show the reference (parent; below) and the query (child; on top) sequences and the inferred TSM processes explaining the differences. The sequence differences are shown in lowercase and the secondary structure is given with the dot-bracket notation. **a** TSM causing a CM with a parallel inversion of the loop sequence (Metazoa, tardigrades). **b**,**c** TSM causing an inversion of the loop sequence only (Bacteria, Actinobacteria; Bacteria, Firmicutes).

To formally test the effect of selection, we studied the ages of the TSM-like changes. In our phylogenetic approach, novel mutations appear as terminal tree branches and split out as sister taxa for the consensus sequence of the main form while older mutations define internal tree branches with descendants inheriting the derived form. Ancestral reconstruction is most robust for sub-terminal nodes, and we recorded the normalized counts of inferred TSMs on the first three levels of phylogenetic trees. We found the TSM-like events to be twice as frequent in the terminal than in sub-terminal (-1, -2) branches, consistent with influence of negative selection (Fig. S3). A mutation being placed in a sub-terminal tree branch does not yet prove it passing evolutionary selection, though it indicates that the underlying mutation is real and the derived form has been independently observed in multiple sequencing experiments.

While the CMs associated with an inversion of the loop sequence were rare, inversions could also occur at a perfectly pairing hairpin sequence and leave no footprint in the adjacent stems. We observed, in total, 69 mutation clusters in the loop sequence consistent with such inversions. The affected loop sequences were typically four nucleotides long and partially reverse complement (85.5% of the cases), minimizing the impact, but we also detected inverted loops of 7-8 nuc in length and containing multiple sequence differences (Fig. 3b,c).

As we searched for CMs consistent with the TSM mechanism, we identified TSM-like mutation patterns that did not originate symmetrically from the opposite stem half but were nevertheless predicted to retain the secondary structure. We focused on three categories: TSM copying sequence from loop to a stem half, TSM within a stem half, and TSM copying a bulge to other stem half. These cases were more common than the TSM-like CMs, totalling 1,788, and statistically enriched in comparison to simulated data (Table 1, Table S1). The cases include an interesting example of an asymmetric from-loop-to-stem mutation first breaking a stem structure and then symmetric TSM fixing the perfect base pairing (Fig. S4).

### Causes and consequences of compensatory mutations

The substitutions patterns of mutations consistent with the two-step CM mechanism show the signal expected from RNA sequences and tend to restore the Watson-Crick pairing of G-U or U-G base pairs (Fig. S5). To allow better comparison to published numbers, we computed similar statistics for instantaneous CMs. For that, we identified loop stems where the parent and child nodes had nonidentical stem sequence but both showed perfect base-pairing (see Methods). We found 68,533 such cases in terminal nodes covering 85,993 CMs, as 21.0% of the stems contained more than one CM (Fig. S6). The numbers of instantaneous and two-step CMs are in line with earlier studies, although their ratio is higher than in the evolutionary models (3; 6). While 78.5% of the identified two-step CMs affected G-U or U-G base pairs and restored Watson-Crick pairing, 61.0% of instantaneous CMs were of type of A-T *↔* G-C or C-G *↔* T-A and could thus occur via a G-U and U-G intermediate (Fig. S5). The fact that a greater proportion of two-step CMs involve G-U/U-G pairs likely reflects their benign impact on the stem structure, enabling higher frequencies in the populations and increased chances of being observed (22; 23).

The high numbers of instantaneous CMs found underlines the modest numbers of two-step cases. On the other hand, the rarity of changes within the loop sequence and the high reverse complementarity of the observed cases of loop inversions is consistent with the enrichment of four nucleotide patterns giving stable RNA structures (20). To understand the effect of this, we computed the frequencies of different rRNA hairpin loops and their reverse complements in our data. We predicted secondary structures to all sequences, and extracted in total 4,874,366 hairpin loop sequences from non-root internal nodes with a terminal node child (see Methods). As expected, we observed enrichment of hairpin sequences belonging to GNRA, UNCG and CUUG families known to form exceptionally stable hairpins (24). The forward sequences for the three classes were, respectively, 36, 66 and 8 fold enriched in comparison to their reverse complements, the most common hairpin sequence GAAA being 99 times more frequent than its opposite UUUC (Table S3). A sequence giving an exceptionally stable loop structure is unlikely to be inverted (Fig. 1f), but it could promote intra-strand template switching (Fig. 1c) by bending the single-strand DNA into a loop conformation. To test this, we computed the frequencies of motifs associated with stable hairpin loop structures and loop lengths within hairpins containing instantaneous CMs, and compared those to background frequencies computed from all hairpins of non-root ancestral sequences. We observed that four-nucleotides-long loops, which represent the most stable rRNA and rDNA loop lengths (25), were strongly enriched among the instantaneous CMs (p-value*<*0.00001, one-proportions z-test; Table 2). Of the most frequent RNA loop motifs, GNRA and CUUG were enriched in our data but UNCG was not (Table 2). Interestingly, GNRA and CUUG are exceptionally stable both in DNA and RNA, whereas UNCG is stable only in RNA (26).

**Table 2.** Frequency of loop sequences and lengths in terminal nodes.

| Motif | Frequency in hairpins |  | p-value |
| --- | --- | --- | --- |
|  | with instant. CM | in full data |  |
| GNRA | 33.98% | 17.90% | 0.000 |
| TNCG | 5.58% | 5.81% | 0.006 |
| CTTG | 2.43% | 0.01% | 0.000 |
| Loop length |  |  |  |
| 3 | 3.52% | 6.11% | 0.000 |
| 4 | 56.01% | 42.13% | 0.000 |
| 5 | 11.70% | 16.79% | 0.000 |
| 6 | 5.22% | 9.88% | 0.000 |
| 7 | 9.57% | 7.56% | 0.000 |
| 8 | 8.93% | 9.47% | 0.000 |

## Discussion

We found footprints of template switch mutations (TSMs) (7; 8) in reconstructed history of RNA sequences and propose the mechanism as the explanation for the origin of perfect stem structures and their evolution through compensatory changes (6; 27). We saw nearly irrefutable evidence for the involvement of template switching in creation of novel stem structures by long sequence insertions. Especially microRNAs, which are both evolutionarily young and structurally extremely simple, provide optimal circumstances for the TSM mechanism to operate. In structurally complex rRNA molecules perfect Watson-Crick base-pair regions tend to be very short (28) but, consistent with our hypothesis, we identified compensatory mutations (CMs) associated with nearby parallel sequence changes explainable by a single TSM event. However, TSM-like events altering the loop sequence, known to be highly conserved (21), were rare and only patterns explaining multiple CMs in the stems were statistically enriched in comparison to simulated data. Although very rare, the presence of complete inversions of the loop sequence suggests that at least some of the TSMs have occurred via the inter-strand mechanism capable of turning the sequence in place. Interestingly, we also observed asymmetric TSM-patterns that retain the RNA secondary structure with sequence coming from outside the opposite stem. These demonstrate the flexibility of RNA sequences and suggest an elegant explanation for the complex mutation patterns, including multi-base changes and jumps in the sequence space observed in evolving RNA sequences (29).

Our study suggests that the TSM mechanism has both a constructive and a destructive role in the evolution of rRNA sequences and structures. The constructive role is supported by the enrichment of structurally exceptionally stable motifs in hairpins associated with instantaneous CMs. Formation of a loop structure is defined by the underlying sequence and 1-2 nucleotides adjacent to the loop region (30; 31). Existing rRNA hairpin loop structures may trigger intra-strand TSMs with the nascent strand forming a loop to prime DNA replication, the sequence for the opposite stem functioning as the template. Crucially, such a mechanism fixes the stem sequence with CMs while retaining the secondary structure and the loop sequence. Similar hairpin loop-priming has been described for inverted repeat-containing oligonucleotides replicated by DNA polymerase I of *Escherichia coli* (32), and replication models of parvovirus (single-strand DNA virus) (33) and poxvirus (double-strand DNA virus) (34). The destructive potential of TSMs is represented by the novel loops and radical changes of the secondary structure. Although they are the ultimate source of novelties and enable the evolution, in an established molecule such as rRNA they tend to be deleterious and radical changes were more frequently observed in terminal branches.

Interestingly, our analyses suggest a solution for another unexplained feature of RNA sequences, the asymmetry of frequencies of G-C and C-G pairs, and of U-A and A-U pairs in the stem structures, the former of each couple appearing at a higher frequency (3). The enrichment of G-C and U-A pairs could be explained by TSMs if the direction of the switch jump and the strand of the switch were conserved. One scenario conserving the direction of switch jump would be the intra-strand TSM events: in those, point ➁ is always left of point ➀ (cf. Fig. 1c vs. Fig. 1d,e) and the base pairing is thus corrected according to the 5’ base. Given this (Fig. 4), the possible two-step CMs involving G-U pairs would always lead to either G-C or U-A base pairing:

**Fig. 4.**
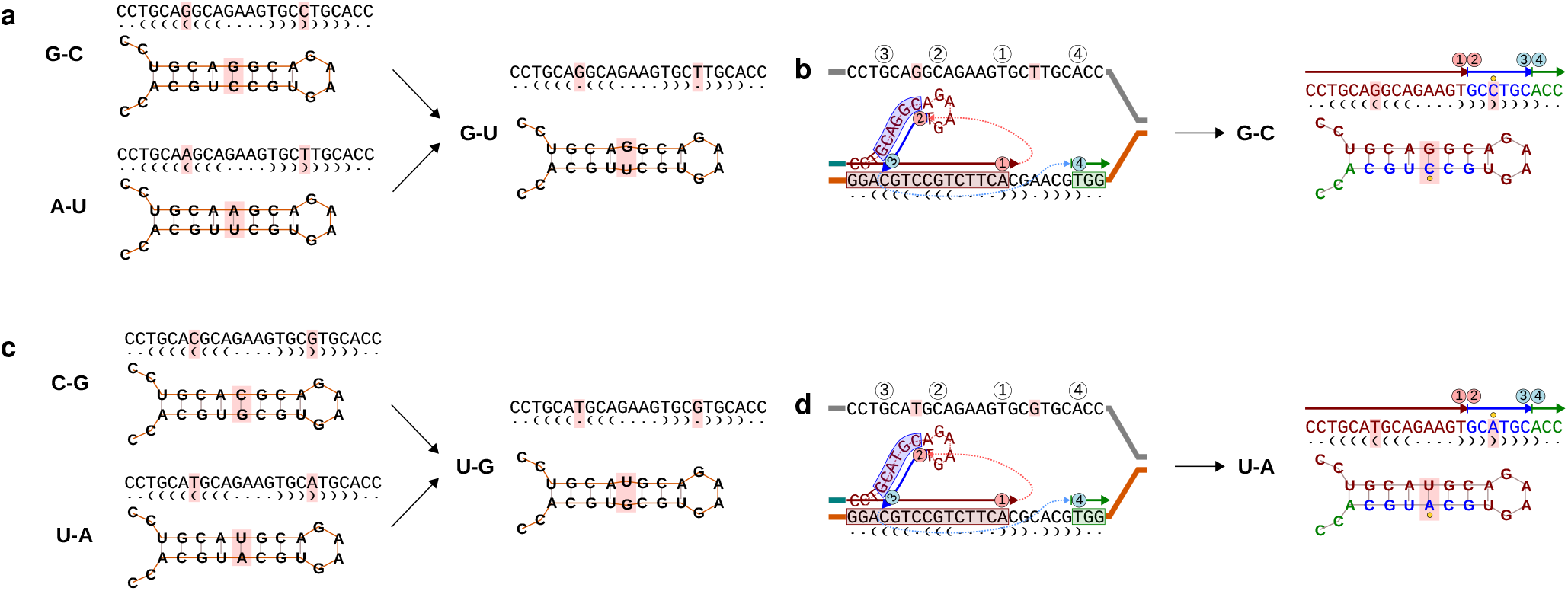
Intra-strand TSMs can explain elevated frequencies of G-C and U-A base pairs in RNA stem regions. **a** A transition converts G-C and A-U pairs to a noncanonical G-U pair. **b** An intra-strand TSM converts the G-U pair to a canonical G-C pair. **c** A transition converts C-G and U-A pairs to a noncanonical U-G pair. **d** An intra-strand TSM converts the U-G pair to a canonical U-A pair.

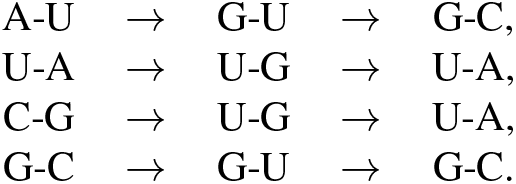

Plausible mechanisms conserving the strand of TSM events could be the coordination of transcription and replication to the same direction, or mutations resulting from clashes of the two systems (35; 36). On the other hand, in the bacterium *E. coli*, template switching has been shown to occur at a much higher frequency on the leading strand than on the lagging strand (37; 38) such that the relative positioning of the replication origin and the rDNA gene cluster could also lead to a bias on the strand of TSM events.

The high numbers of instantaneous CMs observed demonstrate that our data and methodology were adequate for the identification of mutation patterns in ancestral sequence histories. We suspect that the rather modest numbers of positive cases of TSM are, at least partially, explained by the technical challenges of working with data from genes that are both multi-copy and RNA-coding. Ribosomal DNA is known to be a challenging target for phylogenetic analyses (39; 40). More importantly though, isolation of single rDNA genes has so far been difficult and most studied sequences probably represent a consensus formed of multiple, slightly different rDNA copies. In sequence assembly, reads originating from different copies are piled up and, at each position, a call is made among the possible alternatives: given the random sampling of reads across the gene, even the consensus call is volatile and may result in a mixture of different rDNA copies not existing in the nature. Stochasticity of the sequence assembly and true recombinations within rDNA clusters create hybrid sequences, affecting the ancestral state reconstruction and all our down-stream analyses. The situation is improving, however, and with high-throughput long-read DNA sequencing the true content of even complex rDNA gene clusters can be resolved (41). With reads covering full rDNA gene copies it will be possible to study the true variation among the rRNA molecules and distinguish even somatic mutations.

On the other hand, the low numbers of two-step CMs is consistent with the mutation mechanism of rRNA genes and selection keeping the initial mutations at low frequency (3). In multi-copy genes, a mutation initially affects only one gene copy and may then spread within the gene cluster through non-homologous recombination and gene conversion. The latter may refine the mutation and only copy the selectively advantageous parts of the mutation: for example, from a large, abrupt TSM that fixes the base pairing but also breaks the nearby structure, the recombination may pick the beneficial changes and combine them with a functionally working copy. Such a copy can rise in frequency and eventually show up in consensus-based sequences, appearing as an instantaneous compensatory change. Related to this, the final factor reducing the number of positive cases comes from the structure of rRNA molecule and our focus on TSM-patterns of at least six bases in length. While novel hairpin loops of tens of bases in length (cf. Fig. 2a) are practically impossible under the classical mutation processes, the existing perfect Watson-Crick base-pair regions in rRNA genes are usually very short, only a few bases long (28). Due to the low complexity of a four-nucleotide alphabet, a perfect reverse-complement match could be found for nearly any short sequence and the hits have little statistical power (19). Rather counterintuitively, the largest TSM-like pattern that we observed, up to 59 bp in size, were of little interest as they caused major structural changes, altering the loop location or even breaking the hairpin structure, and failed in quality control.

Overall, our analyses demonstrate that a TSM-like mechanism can create novel hairpin stem structures with a one-off mutation event. The mechanism’s role in the maintenance of short RNA stems with compensatory changes is less clear, the strongest evidence being circumstantial, but it does provide elegant explanations for several open questions in RNA evolution. We did not make assumptions about the biological mechanism(s) behind the observed mutations but focused on local events within stem loop structures. Whatever the mechanism, it could allow a longer distance between the source and target regions (see e.g. (42)) and thus have a greater role in the RNA evolution than suggested by our study. While spatially more distant copying events would not create hairpins, they would still generate multiple parallel changes – inexplicable by classical models - and may be crucial for RNA tertiary structures. Interestingly, multi-copy gene clusters may be affected by non-replication-related mutation mechanisms (43) with the potential of creating similar mutation patterns. This has no impact on our findings, however, and the explanation for parallel compensatory substitutions and perfect stem structures would still be copying DNA from another template.

## Methods

### Data collection and preprocessing

The ribosomal RNA sequences were downloaded from Silva metagenomic database v. 138 (http://www.arb-silva.de (16)) and Rfam database v. 14.1 (ftp://ftp.ebi.ac.uk/pub/databases/Rfam (18)) (RNA families RF00177, RF01959, RF01960, RF02540, RF02541 and RF02543). Taxonomies for the downloaded RNA sequences were received from Silva database (v. 138) and NCBI databases (ftp://ftp.ncbi.nih.gov/pub/taxonomy/accession2taxid/ for taxonomy identifiers and ftp://ftp.ncbi.nlm.nih.gov/pub/taxonomy/newtaxdump/ for scientific names; downloaded on 22 March 2020). Information on the RNA families were parsed from the Rfam seed file for v. 14.1 (ftp://ftp.ebi.ac.uk/pub/databases/Rfam/14.1/Rfam.seed.gz). A custom python script was used to remove sequences with degenerate IUPAC symbols and to replace U’s with T’s. Identical sequences were removed using the cd-hit-est tool of the CD-HIT package v. 4.7 (44; 45) with wordsize 11 and retaining the longest representative. Sequences were then clustered using the cd-hit-est applying wordsize 8, 90% sequence identity and 75% minimum alignment coverage for the longest and shortest sequences (parameters -n, -c, -aL and -aS respectively), and divided into the clusters of minimum ten sequences using the make multi seq.pl script. Within clusters, the sequences were compared pairwise and if two sequences did not have at least two mismatches within a 20 bp window, latter of the pair was discarded. Known microRNA loci were downloaded from Ensembl v. 103 (46) using the Ensembl REST API (47).

### Ancestral sequence history and inference of TSMs

Sequence clusters were aligned with MAFFT (v. 7.310; FFT-NS-i, 1,000 iterations) (48), and then trimmed with TrimAl (49) and ‘automated’ mode. Alignments shorter than 200 columns were discarded. Maximum likelihood (ML) trees were computed with IQ-TREE (v. 1.6.1) (50) using automated model selection (51) and tree finder, and resulting trees were midpoint-rooted using ete3 python library (52). Sequences within each cluster were realigned and ancestral sequences inferred according to ML trees with PAGAN2 (53). Phylogenetic trees were traversed with a custom python script applying ete3 library (52): each non-root parent node was compared to its child node (query node) using the FPA tool (see Table S4 for details) and mutation patterns consistent with TSM events of at least six nucleotides in length were recorded.

Instantaneous CMs differ from two-step CMs potentially explainable by TSMs and have the initial mutation and the restoring mutation present in opposite stems of the same sequence. They were identified by screening parent-descendant pairs in each tree and locating loops that i) were in identical positions in both sequences, ii) formed perfect Watson-Crick base pairing, and iii) differed in sequence. Uncertain IUPAC characters were not allowed in reference hairpin sequences. microRNA data were analyzed similarly with ancestral sequence histories inferred on the PAGAN2 alignment and the original EPO phylogeny.

### Sequence simulations

Using the dawg software(54), five replicates were simulated of each sequence cluster according to the original inferred tree and using one randomly selected terminal sequence as a root sequence. The base frequencies, substitution rates and gamma shape alpha for the simulation model were calculated using IQ-TREE (v. 1.6.1). The branch lengths for full-length sequences were calculated under GTR model using Treetime python package (v.0.8.3.1). The gapModel and indel rates were inferred with the lambda.pl script (dawg). The separate rates for insertions and deletions were obtained by diving the indel rate by two.

### Quality control for inferred TSMs

The secondary structures were predicted using RNAfold (55) with default parameters. High quality regions were inferred separately for each internal node by comparing the sequence and secondary structure of the target node to those of its child nodes, and the sequence of the target node to that of its own parent; before the comparison, homologous uncertainties were unified at sites with overlapping base (IUPAC symbols) ranges. Sites with identical base or secondary structure characters were recorded. Using a 10-base sliding window, windows with i) at least nine base identities in either child sequence, ii) at least six identities in the corresponding child secondary structure, and iii) nine base identities in the ancestor sequence passed the quality criteria.

In inferred TSM events, the target region at a child node differs from the homologous sequence at its parent node and the differences can be attributed to the TSM mechanism copying sequence from the nearby source region (at the parent) in reverse-complement manner. In two-step mutations, the TSM source and target regions had to be perfectly reverse-complement, locate in high-quality regions and the source site had to be identical between the sequences if it did not overlap with the target region; if the child node was an internal node, the source and target regions had to be inherited by at least one of its own child nodes in identical form. In the case of instantaneous CMs, the loop had to be a part of high quality regions and identically located in the parent and child nodes; if the child node was an internal node, the loop sequence had to be inherited by at least one of its own child nodes in identical form. For both CM types, only one of multiple fully overlapping cases was counted. The sum of internal node bases within high quality regions was considered the effective length of the data. Correction factors for the simulation replicates were calculated by dividing the effective length of the empirical data by that of the simulated data.

Sequences of all solved structures in PDB were downloaded (ftp://ftp.wwpdb.org/pub/pdb/derived_data/pdb_seqres.txt.gz; downloaded on 28 October 2020). The sequences for rRNA were extracted based on the title (982 sequences in total), a BLAST database was created, and all terminal sequences were screened against it using blastn (v. 2.6.0+). Hits with e-value 0.00 and sequence identity of at least 95% for a sequence containing a TSM hit were selected using awk and python scripts. The structures for these were downloaded from PDB (ftp://ftp.ebi.ac.uk/pub/databases/pdb/compatible/pdb_bundle), and the affected hairpin chain was extracted. Predicted and pdb structures were visualized using RNApdbee 2.0 (56) (http://rnapdbee.cs.put.poznan.pl) with default settings with the exception of the visualization setting ‘PseudoViewer-based procedure’.

### TSMs’ impact on secondary structures

A *compensating mutation* was attributed to a TSM event if i) either TSM source or target sequence overlapped with a sequence difference between the query and its parent, ii) source and target were in the separate halves of the same stem, iii) source and target were equally distant from the loop region, and iv) the mutation corrects an imperfect Watson-Crick base pair into a perfect one. Internal loops or bulges were not taken into account. A *loop inversion* was attributed to a TSM event if i) source and target overlapped with both stem halves of a loop and ii) the length of the loop was maintained. A TSM was considered *asymmetric* if source and target were not equally far from the loop region. A stem was considered *extended* by a TSM event if source and target were on different stem halves and one stem half had gained an insertion.

Statistical significance of observed TSM patterns was studied using independent sample t-test and comparing the numbers of cases of each category in empirical and simulated data separately for terminal and internal nodes. Frequencies were calculated for each category independently in each cluster. The counts for simulated data were multiplied by a correction factor.

### Calculation of TSM and base frequencies

TSM-patterns associated with CMs, loop inversions or insertions in a stem half were counted at each tree level and the counts were divided by the total number of branches of that tree level in the affected trees. The identification of the branch level was done with post-order tree traversal, assigning each branch to the lowest possible level. Hairpin loop sequences in secondary structures of internal nodes were extracted and counted. The sequence and structural quality of the hairpins were confirmed as described above; in addition, the closing pair had to form a perfect Watson-Crick base-pair and uncertain IUPAC characters were not allowed in the hairpin sequence. The frequencies of different loop lengths and loop sequence motifs were compared between hairpins with instantaneous CMs and all hairpins. The significance of the differences was studied using two-sided one proportion z-test. Mutation types in the stems (base pair in parent vs. child) were counted separately for instantaneous and two-step CMs, and the counts were visualized as a heatmap using the R package ggplot2.

### Script availability

The scripts used in the study are available at Github (https://github.com/helimonttinen/TSM_project).

## Acknowledgements

This study was enabled by the Academy of Finland grant #322681.

## References

[1] PB Moore, Structural Motifs in RNA. Annual Review of Biochemistry 68, 287–300 (1999).

[2] N Leontis, E Westhof, Modeling RNA Molecules, eds. N Leontis, e Westhof. (Springer Berlin Heidelberg, Berlin, Heidelberg) Vol. 27, pp. 5–17 (2012).

[3] PG Higgs, RNA secondary structure: physical and computational aspects. Quarterly Reviews of Biophysics 33, 199–253 (2000).

[4] Woese, C R and Pace, N R, Probing RNA Structure, Function, and History by Comparative Analysis, ed. R. F. Gesteland and J F Atkins. (Cold Spring Harbor Laboratory Press), pp. 91–117 (1993).

[5] RR Gutell, N Larsen, CR Woese, Lessons from an evolving rRNA: 16S and 23S rRNA structures from a comparative perspective. Microbiology and Molecular Biology Reviews 58, 10–26 (1994).

[6] ER Tillier, RA Collins, High apparent rate of simultaneous compensatory base-pair substitutions in ribosomal RNA. Genetics 148, 1993–2002 (1998).

[7] LS Ripley, Model for the participation of quasi-palindromic DNA sequences in frameshift mutation. Proceedings of the National Academy of Sciences 79, 4128–4132 (1982).

[8] BE Dutra, S. Lovett, Cis and Trans-acting Effects on a Mutational Hotspot Involving a Replication Template Switch. Journal of Molecular Biology 356, 300–311 (2006).

[9] A Löytynoja, N Goldman, Short template switch events explain mutation clusters in the human genome. Genome Research 27, 1039–1049 (2017).

[10] ED Ladoukakis, A Eyre-Walker, The excess of small inverted repeats in prokaryotes. J. Mol. Evol. 67, 291–300 (2008).

[11] B Lavi, E Levy Karin, T Pupko, E Hazkani-Covo, The Prevalence and Evolutionary Conservation of Inverted Repeats in Proteobacteria. Genome Biology and Evolution 10, 918–927 (2018).

[12] RR Sinden, RD Wells, DNA structure, mutations, and human genetic disease. Curr. Opin. Biotechnol. 3, 612–622 (1992).

[13] L Costantino, et al., Break-Induced Replication Repair of Damaged Forks Induces Genomic Duplications in Human Cells. Science 343, 88–91 (2014).

[14] CMB Carvalho, JR Lupski, Mechanisms underlying structural variant formation in genomic disorders. Nature Reviews Genetics 17, 224–238 (2016).

[15] ED Ladoukakis, A Eyre-Walker, Searching for sequence directed mutagenesis in eukaryotes. J. Mol. Evol. 64, 1–3 (2007).

[16] C Quast, et al., The SILVA ribosomal RNA gene database project: improved data processing and web-based tools. Nucleic Acids Research 41, D590–596 (2013).

[17] JF Elder, BJ Turner, Concerted Evolution of Repetitive DNA Sequences in Eukaryotes. The Quarterly Review of Biology 70, 297–320 (1995).

[18] I Kalvari, et al., Non-Coding RNA Analysis Using the Rfam Database. Current Protocols in Bioinformatics 62, e51 (2018).

[19] CR Walker, A Scally, N De Maio, N Goldman, Shortrange template switching in great ape genomes explored using pair hidden markov models. PLOS Genetics 17, 1–30 (2021).

[20] G Varani, Exceptionally Stable Nucleic Acid Hairpins. Annual Review of Biophysics and Biomolecular Structure 24, 379–404 (1995).

[21] S Smit, J Widmann, R Knight, Evolutionary rates vary among rRNA structural elements. Nucleic Acids Research 35, 3339–3354 (2007).

[22] W Stephan, The rate of compensatory evolution. Genetics 144, 419—426 (1996).

[23] PG Higgs, Compensatory neutral mutations and the evolution of RNA. Genetica 102-103, 91–101 (1998).

[24] HA Heus, A Pardi, Structural features that give rise to the unusual stability of RNA hairpins containing GNRA loops. Science 253, 191–194 (1991).

[25] DR Groebe, OC Uhlenbeck, Characterization of RNA hairpin loop stability. Nucleic Acids Research 16, 11725–11735 (1988).

[26] VP Antao, SY Lai, I Tinoco, A thermodynamic study of unusually stable RNA and DNA hairpins. Nucleic Acids Research 19, 5901–5905 (1991).

[27] MT Dixon, DM Hillis, Ribosomal RNA secondary structure: compensatory mutations and implications for phylogenetic analysis. Molecular Biology and Evolution 10, 256–267 (1993).

[28] J Wolters, The nature of preferred hairpin structures in 16S-like rRNA variable regions. Nucleic Acids Research 20, 1843–1850 (1992).

[29] MV Meer, AS Kondrashov, Y Artzy-Randrup, FA Kondrashov, Compensatory evolution in mitochondrial tRNAs navigates valleys of low fitness. Nature 464, 279–282 (2010).

[30] MT Woodside, et al., Nanomechanical measurements of the sequence-dependent folding landscapes of single nucleic acid hairpins. Proceedings of the National Academy of Sciences 103, 6190–6195 (2006).

[31] G Bonnet, O Krichevsky, A Libchaber, Kinetics of conformational fluctuations in DNA hairpin-loops. Proceedings of the National Academy of Sciences 95, 8602–8606 (1998).

[32] E Uhlmann, An alternative approach in gene synthesis: use of long selfpriming oligodeoxynucleotides for the construction of double-stranded DNA. Gene 71, 29–40 (1988).

[33] SF Cotmore, P Tattersall, Parvovirus diversity and DNA damage responses. Cold Spring Harbor Perspectives in Biology 5 (2013).

[34] B Moss, Poxvirus DNA replication. Cold Spring Harbor Perspectives in Biology 5 (2013).

[35] S Hamperl, MJ Bocek, JC Saldivar, T Swigut, KA Cimprich, Transcription-Replication conflict orientation modulates R-Loop levels and activates distinct DNA damage responses. Cell 170, 774–786.e19 (2017).

[36] YH Chen, et al., Transcription shapes DNA replication initiation and termination in human cells. Nat. Struct. Mol. Biol. 26, 67–77 (2019).

[37] WA Rosche, TQ Trinh, RR Sinden, Leading strand specific spontaneous mutation corrects a quasipalindrome by an intermolecular strand switch mechanism. J. Mol. Biol. 269, 176–187 (1997).

[38] T Seier, et al., Insights into mutagenesis using escherichia coli chromosomal lacz strains that enable detection of a wide spectrum of mutational events. Genetics 188, 247–262 (2011).

[39] H Letsch, K Kjer, Potential pitfalls of modelling ribosomal rna data in phylogenetic tree reconstruction: Evidence from case studies in the metazoa. BMC evolutionary biology 11, 146 (2011).

[40] TH Eickbush, DG Eickbush, Finely Orchestrated Movements: Evolution of the Ribosomal RNA Genes. Genetics 175, 477–485 (2007).

[41] S Nurk, et al., The complete sequence of a human genome. (2021).

[42] WM Hicks, M Kim, JE Haber, Increased mutagenesis and unique mutation signature associated with mitotic gene conversion. Science 329, 82–85 (2010).

[43] D Liao, Concerted Evolution: Molecular Mechanism and Biological Implications. The American Journal of Human Genetics 64, 24–30 (1999).

[44] L Fu, B Niu, Z Zhu, S Wu, W Li, CD-HIT: accelerated for clustering the next-generation sequencing data. Bioinformatics 28, 3150–3152 (2012).

[45] W Li, A Godzik, Cd-hit: a fast program for clustering and comparing large sets of protein or nucleotide sequences. Bioinformatics 22, 1658–1659 (2006).

[46] AD Yates, et al., Ensembl 2020. Nucleic Acids Res. 48, D682–D688 (2020).

[47] A Yates, et al., The ensembl REST API: Ensembl data for any language. Bioinformatics 31, 143–145 (2015).

[48] K Katoh, DM Standley, MAFFT multiple sequence alignment software version 7: improvements in performance and usability. Molecular Biology and Evolution 30, 772–780 (2013).

[49] S Capella-Gutiérrez, JM Silla-Martínez, T Gabaldón, trimAl: a tool for automated alignment trimming in largescale phylogenetic analyses. Bioinformatics 25, 1972–1973 (2009).

[50] LT Nguyen, HA Schmidt, A von Haeseler, BQ Minh, IQ-TREE: a fast and effective stochastic algorithm for estimating maximum-likelihood phylogenies. Molecular Biology and Evolution 32, 268–274 (2015).

[51] S Kalyaanamoorthy, BQ Minh, TKF Wong, A von Haeseler, LS Jermiin, ModelFinder: fast model selection for accurate phylogenetic estimates. Nature Methods 14, 587–589 (2017).

[52] J Huerta-Cepas, F Serra, P Bork, ETE 3: Reconstruction, Analysis, and Visualization of Phylogenomic Data. Molecular Biology and Evolution 33, 1635–1638 (2016).

[53] A Löytynoja, AJ Vilella, N Goldman, Accurate extension of multiple sequence alignments using a phylogeny-aware graph algorithm. Bioinformatics 28, 1684–1691 (2012).

[54] RA Cartwright, DNA assembly with gaps (dawg): simulating sequence evolution. Bioinformatics 21 Suppl 3, iii31–8 (2005).

[55] R Lorenz, et al., ViennaRNA Package 2.0. Algorithms for Molecular Biology 6, 26 (2011).

[56] M Antczak, et al., RNApdbee—a webserver to derive secondary structures from pdb files of knotted and unknotted RNAs. Nucleic Acids Research 42, W368–W372 (2014).

